# Understanding and predicting ciprofloxacin minimum inhibitory concentration in *Escherichia coli* with machine learning

**DOI:** 10.1101/806760

**Authors:** Bálint Ármin Pataki, Sébastien Matamoros, Boas C.L. van der Putten, Daniel Remondini, Enrico Giampieri, Derya Aytan-Aktug, COMPARE ML-AMR group, Rene S. Hendriksen, Ole Lund, István Csabai, Constance Schultsz

**Author notes:** see the full list of the COMPARE ML-AMR group members in acknowledgements. corresponding author: Bálint Ármin Pataki. **1.5 Repositories:** Our work does not contain new sequence data.

## Abstract

2.

A possible way to tackle the crisis of antimicrobial resistance development is a strict policy when prescribing antibiotics. Thus, it is important that prescriptions are based on antimicrobial susceptibility data to ensure effective treatment outcomes. The increasing availability of next-generation sequencing (NGS), bacterial whole genome sequencing (WGS) can facilitate a more reliable and faster alternative to traditional phenotyping for the detection and surveillance of AMR.

This work proposes a machine learning approach that can predict the minimum inhibitory concentration (MIC) for a given antibiotic, here ciprofloxacin, on the basis of both genome-wide mutation profiles and profiles of acquired antimicrobial resistance genes (ARG). We analysed 704 *Escherichia coli* genomes combined with their respective MIC measurements for ciprofloxacin originating from different countries. The four most important predictors found by the model, mutations in *gyrA* residues Ser83 and Asp87, a mutation in *parC* residue Ser80 and presence of any *qnrS* gene, have been experimentally validated before. Using only these four predictors in a linear regression model, 65% and 92% of the test samples’ MIC were correctly predicted within a two- and a four-fold dilution range, respectively. The presented work goes further than the typical predictions that use machine learning as a black box model concept. The recent progress in WGS technology in combination with machine learning analysis approaches indicates that in the near future WGS of bacteria might become cheaper and faster than a MIC measurement.

**Impact statement:** Whole genome sequencing has become the standard approach to study molecular epidemiology of bacteria. However, the application of WGS in the clinical microbiology laboratory as part of individual patient diagnostics still requires significant steps forward, in particular with respect to prediction of antibiotic susceptibility based on DNA sequence. Whilst the majority of studies of prediction of susceptibility have used a binary outcome (susceptible/resistant), a quantitative prediction of susceptibility, such as the MIC, will allow for earlier detection of trends in increasing resistance as well as the flexibility to follow potential adjustments in definitions of susceptible (wild type) and resistant (non-wild type) categories (breakpoints/ epidemiological cut-off values).

**Data summary:** In this study, 704 *E. coli* genomes combined with MIC measurement for ciprofloxacin were analysed (24). Paired-end sequencing was performed on all isolates and the results were stored in FASTQ format. The isolates originated from five countries, Denmark, Italy, USA, UK, and Vietnam. The MIC distribution for these isolates is depicted in Table 1. Out of 704, 266 *E. coli* genomes had no country metadata available and were used as an independent test set. All data were deposited in the AMR Data Hub (24) which consists of raw sequencing data, ciprofloxacin minimum inhibitory concentrations, and additional metadata such as the origin of the samples.

TABLE 1
The collected and used data in the analysis grouped by country and MIC values.

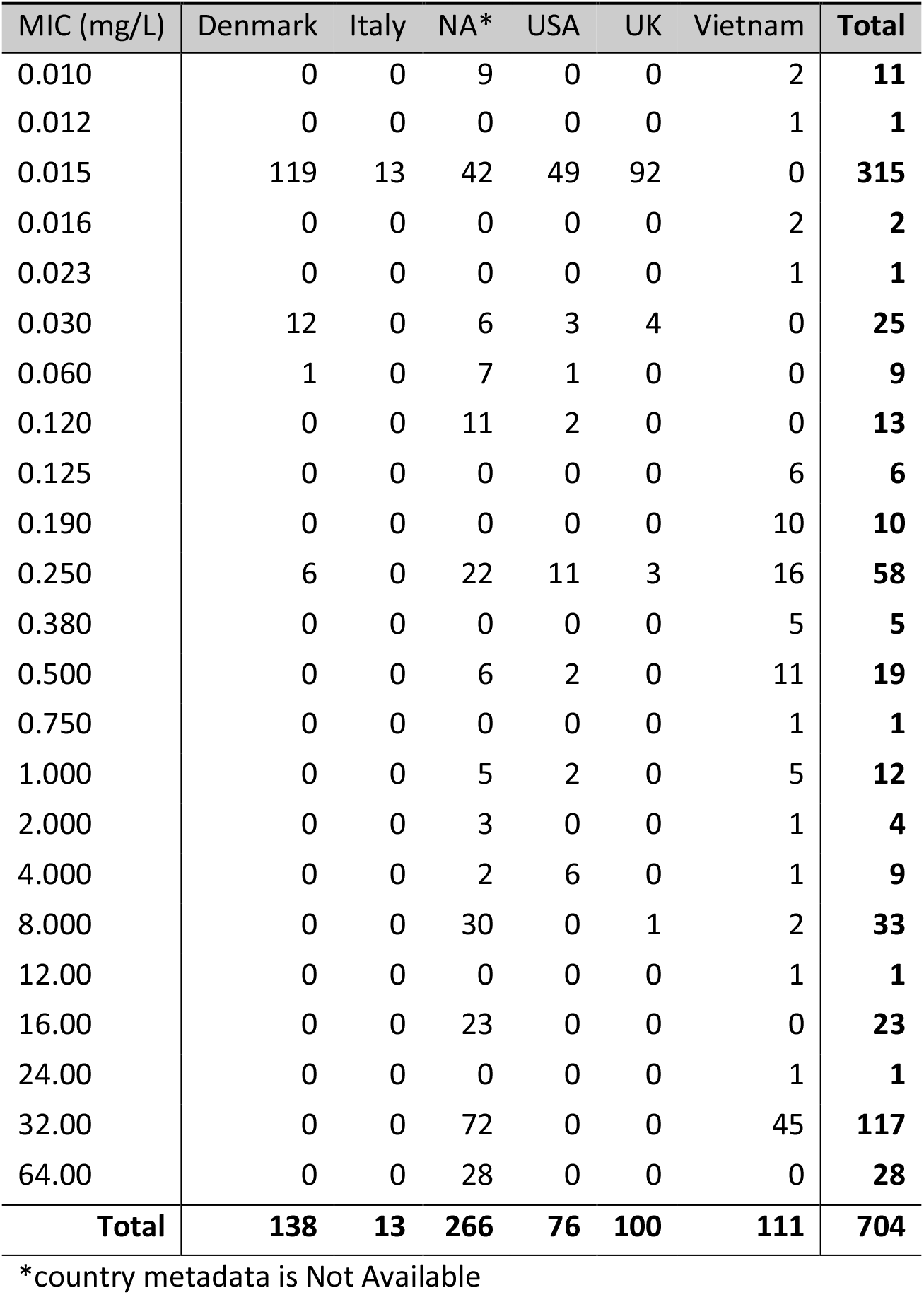

**TABLE 1** The collected and used data in the analysis grouped by country and MIC values.
| MIC (mg/L) | Denmark | Italy | NA* | USA | UK | Vietnam | Total |
| --- | --- | --- | --- | --- | --- | --- | --- |
| 0.010 | 0 | 0 | 9 | 0 | 0 | 2 | **11** |
| 0.012 | 0 | 0 | 0 | 0 | 0 | 1 | **1** |
| 0.015 | 119 | 13 | 42 | 49 | 92 | 0 | **315** |
| 0.016 | 0 | 0 | 0 | 0 | 0 | 2 | **2** |
| 0.023 | 0 | 0 | 0 | 0 | 0 | 1 | **1** |
| 0.030 | 12 | 0 | 6 | 3 | 4 | 0 | **25** |
| 0.060 | 1 | 0 | 7 | 1 | 0 | 0 | **9** |
| 0.120 | 0 | 0 | 11 | 2 | 0 | 0 | **13** |
| 0.125 | 0 | 0 | 0 | 0 | 0 | 6 | **6** |
| 0.190 | 0 | 0 | 0 | 0 | 0 | 10 | **10** |
| 0.250 | 6 | 0 | 22 | 11 | 3 | 16 | **58** |
| 0.380 | 0 | 0 | 0 | 0 | 0 | 5 | **5** |
| 0.500 | 0 | 0 | 6 | 2 | 0 | 11 | **19** |
| 0.750 | 0 | 0 | 0 | 0 | 0 | 1 | **1** |
| 1.000 | 0 | 0 | 5 | 2 | 0 | 5 | **12** |
| 2.000 | 0 | 0 | 3 | 0 | 0 | 1 | **4** |
| 4.000 | 0 | 0 | 2 | 6 | 0 | 1 | **9** |
| 8.000 | 0 | 0 | 30 | 0 | 1 | 2 | **33** |
| 12.00 | 0 | 0 | 0 | 0 | 0 | 1 | **1** |
| 16.00 | 0 | 0 | 23 | 0 | 0 | 0 | **23** |
| 24.00 | 0 | 0 | 0 | 0 | 0 | 1 | **1** |
| 32.00 | 0 | 0 | 72 | 0 | 0 | 45 | **117** |
| 64.00 | 0 | 0 | 28 | 0 | 0 | 0 | **28** |
| **Total** | **138** | **13** | **266** | **76** | **100** | **111** | **704** |

\*country metadata is Not Available

Publicly available sequencing data was used from projects PRJEB21131, PRJNA266657, PRJNA292901, PRJNA292904, PRJNA292902, PRJDB7087, PRJEB21880, PRJEB21997, PRJEB14086 and PRJEB16326.

Download and analysis scripts are available at https://github.com/patbaa/AMR_ciprofloxacin. iTOL phylogenetic tree is available at https://itol.embl.de/tree/14511722611491391569485969.

**The authors confirm all supporting data, code and protocols have been provided within the article or through supplementary data files.**

## 5. Introduction

Antibiotics are an essential resource in the control of infectious diseases; they have been a major contributor to the decline of infection-associated mortality and morbidity in the 20th century. However, the recent rise of antimicrobial resistance (AMR) threatens this situation (1). Bacterial AMR is associated with a higher likelihood of therapeutic failure in case of infections. Accurate and fast prediction of AMR in bacteria is needed to select the optimal therapy.

With the increasing availability of next-generation sequencing (NGS), bacterial whole genome sequencing (WGS) is becoming a feasible alternative to traditional phenotyping for the detection and surveillance of AMR (2), (3), (4). However, data analysis remains the weak point in this approach; fast and scalable methods are required to transform the ever-growing amount of genomic data into actionable clinical or epidemiological information (5). Machine learning is a promising approach for this kind of data analysis.

AMR can be predicted in numerous ways. In addition to classic and highly standardized phenotypic testing of resistance, several methods of resistance prediction have been developed. Most novel methods use a genetic or genomic approach, although transcriptomic approaches have been investigated as well (6), (7), (8). An important factor in the choice of the resistance prediction method is the microorganism under study. For example, the CRyPTIC consortium managed to predict resistance to four first-line drugs in *Mycobacterium tuberculosis,* using only known mutations extracted from WGS (9). However, *M. tuberculosis* displays little-to-no horizontal gene transfer and low genomic evolution rate (10), which makes it feasible to predict resistance only on the basis of known mutations (11). For other bacteria, more advanced analysis methods such as machine learning need to be applied to allow for accurate prediction.

Machine learning has been applied to predict resistance from WGS data in several settings. To date, these methods have been restricted mostly to assign bacteria to binary categories, i.e. susceptible or non-susceptible (12), (13), (14), (8), (15), (16), (17). However, clinical breakpoints used to define susceptible and non-susceptible categories can change and such binary categories do not allow following more trends in subtle changes in susceptibility in time. MIC measures offer an adequate resolution to follow if susceptibility is changing in a population, which is useful for epidemiological purposes. Therefore, a resistance prediction method would preferably output a continuous estimate of resistance similar to MIC, instead of binary classification (S/R), as a number of studies already proposed (18), (19), (20), (21).

Additional issues should be considered when developing a reliable and useful prediction model. First, genotypes are often geographically clustered (22). This implies that if a prediction model is trained on data from one country, this model might not be generalizable to data from another country. Data from multiple countries are thus needed. Secondly, complex combinations of chromosomal point mutations and acquired resistance genes influence antimicrobial resistance. Therefore, different data types need to be combined to obtain a biologically relevant set of input data. Lastly, while machine learning is able to analyse highly complex patterns of features, the model would preferably output generally understandable data. K-mer profiles have been used to predict resistance, but these can be difficult to interpret (19) (20).

In this study, we focus on predicting a quantitative measure of ciprofloxacin resistance (MIC) for a geographically diverse population of *E. coli* using machine learning. We chose to study ciprofloxacin resistance in *E. coli* because of three reasons:

1. Ciprofloxacin resistance in *E. coli* has been studied intensively
2. Ciprofloxacin resistance in *E. coli* can be caused by a range of different chromosomal and plasmid-mediated mechanisms (23)
3. Ciprofloxacin is commonly used in the treatment of *E. coli* infections across the globe.

In our selection of machine learning models, an important criterion was that high-scoring features could be extracted from the model. This would allow us to explore the reasoning behind each prediction and thus to interpret and understand the model.

## 6. Methods

### 6.1 Data preprocessing

Raw reads were mapped on the ATCC 25922 reference genome (https://www.ncbi.nlm.nih.gov/assembly/GCF_000743255.1) using BWA-MEM v0.7.17 (25) with default settings. Pileup files were generated with bcftools v1.9 (26) with “–min-MQ 50” settings. Single-nucleotide polymorphisms (SNPs) and insertions-deletions (INDELs) were called using bcftools v1.9 with “–ploidity 1−m “ flags. Further filtering was applied via bcftools v1.9 “%QUAL>=50 & DP>=20” flags. Bcftools output data was expressed as either a SNP (value: 1), an INDEL (value: 5) or no mutation (value: 0) per position in the reference genome. Exact numbers are irrelevant, as tree-based methods are not sensitive to the scale. The intention was to differentiate between reference alleles, SNPs and INDELs at a given position. Acquired resistance genes were identified using ResFinder v3.1.0 (27) with a coverage threshold of 60% and an identity threshold of 90% using a database downloaded on 5th Dec. 2018. ResFinder was used with KMA v1.1.4 (28). The ResFinder output data was expressed as presence (value: 1) or absence (value: 0) of resistance genes. The SNP/INDEL data and ResFinder data were subsequently merged which provided a matrix with more than 830,000 columns representing reference genome positions with at least one mutation and 959 columns representing detected resistance genes. Two more binary columns were added manually, which describe if any *qnr* or *qnrS* gene is present in the given genome or not. Grouping together *qnr* and *qnrS* genes such a way was motivated by their proven role in ciprofloxacin resistance (23). In the ResFinder database these genes have many different variants and for the machine learning models it helps to provide aggregated information too. The different specific gene variants may be present only in a few isolates each.

### 6.2 Phylogenetic tree generation

The merged variant call files were converted to a FASTA alignment using vcf2phylip v2.0, retaining positions that were called in at least 50% of isolates (29). The invariant positions were removed from the alignment using snp-sites v2.4.0 (30). The phylogeny was inferred using RAxML v8.2.9 in rapid bootstrap mode (-f a) with 100 bootstraps using a General Time Reversible model with Gamma rate heterogeneity including Lewis ascertainment bias correction (-m ASC_GTRGAMMA) (31). The resulting phylogeny was visualized in iTOL (32).

### 6.3 Metrics

We used the following metrics for the evaluation of the model:

<u>**AUC** - area under the receiver operating characteristics curve</u> - we used the clinical breakpoint for ciprofloxacin, 1 mg/L, based on the Clinical & Laboratory Standards Institute guideline (33) to binarize the samples whether they are resistant or not.
<u>**R^2^ score** - coefficient of determination</u>

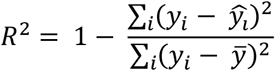

where

- *y_i_* is the true value for sample i,
- *ŷ_l_* is the predicted value for sample i,
- *ȳ* is the mean of the true values.
<u>**R** - Pearson correlation coefficient</u>

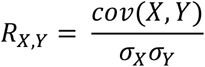

where *cov* is the covariance and *σ* is the standard deviation.
<u>**ME** - major error</u> - when the sample is non-resistant by measurement, but it is predicted to be resistant. Non-resistant and resistant labels are derived from MIC via thresholding.
<u>**VME** - very major error</u> - when the sample is resistant by measurement, but it is predicted to be non-resistant. Non-resistant and resistant labels are derived from MIC via thresholding.
<u>**ACC-2** - accuracy within two-fold dilution</u> - the fraction of the samples with MIC properly predicted within a two-fold dilution. If the measured MIC is x, then the prediction is counted as properly predicted within a two-fold dilution if it falls to the [x/2;2x] interval.
<u>**ACC-4** - accuracy within four-fold dilution</u> - the fraction of the samples with MIC properly predicted within a four-fold dilution. If the measured MIC is x, then the prediction is counted as properly predicted within a four-fold dilution if it falls to the [x/4;4x] interval.
<u>**MAFE** - mean absolute fold error</u> - The mean absolute difference between the log2 values of the prediction and the measurements.

### 6.4 Importance of the validation scheme

Proper validation is a key element in machine learning as most of the models have a large number of parameters. In image recognition, popular convolutional neural networks can have more than 100M parameters (34). This number of parameters is orders of magnitudes larger than the number of pixels of a single image or even the number of the images in the full usual training data set, such as ImageNet (35). Having that many parameters it is possible to memorize the training data without generalizing any knowledge to the test data or for future use.

However, with having a proper validation scheme it is easy to test the generalization power of a model. In many cases simply randomly splitting the samples into two groups to a test and a validation set is enough. If the data set is small, cross-validation is needed, usually, K-fold cross-validation, where the data set is split into K set, each having the same size. Then, the model is trained on using data from K – 1 set and the predictions are made for the one set that was not used in the training process. Repeating the process, K times, predictions can be generated for the whole data set in a way that the model did not see in training time any of the samples for which it is generating predictions. The weights of the model are reset between any two training.

K-fold cross-validation can produce too optimistic results if the samples are clustered. For example, when the data collection is biased, bacterial isolates from one country are predominantly resistant whilst isolates from other countries are predominantly susceptible to an antibiotic. In addition, genetic signatures are often clustered by country (22). Due to such clustering, the model may predict based on the country of origin of the bacterial isolate, which may be correlated with the MIC, on both the training and the validation data sets, but it is not guaranteed that the same will happen in real-life usage later.

#### 6.4.1 Leave-one-country-out validation

Here we propose a more strict and reliable validation method. Instead of randomly splitting the data into K different folds, we split the folds by country. Using this approach, the model is not rewarded if it only learns country-specific attributes. Leave-one-country-out validation was performed during the selection of the most important features in the data set, see Table S1. The random forest model was fitted K = 5 times leaving out one country each time from the training data set. Then the feature importances were summed over each fold resulting in the final feature importance rankings.

### 6.5 Random forest model

For tabular data most often tree-like models perform the best. The random forest model is an ensemble of numerous (usually hundreds of) decision trees. In the training process, each tree is trained separately and each of them uses only a random fraction of the data, which ensures that the decision trees will not be identical. For a new sample, the prediction is the average of the prediction of the trees, or for classification the category that was predicted the most often by the individual trees. This ensemble technique ends up an accurate, robust, scalable model. The prediction error is usually large for each individual tree, but as long as the errors of the trees are uncorrelated, averaging their prediction lowers the final error. Random forest regressor was trained on the whole training dataset with mean squared error criterion, n_estimators = 200 for the feature selection. For the final evaluation mean squared error criterion, min_samples_split = 5, and n_estimators = 100 parameters were used. The random seed was fixed for both model to be 42. Other parameters remained default. Scikit-learn v0.21.2 (36) was used for fitting the model in Python 3.6.5.

#### 6.5.1 Random forest feature importance

For decision trees, the input variables can be ranked by their contribution to the predictions. This score is called feature importance. The importance can be defined in various ways; the used scikit-learn v0.21.2 (36) implementation calculates the mean decrease impurity averaged over all the trees in the forest (37) (38). In this approach, the identification of the most important predictors becomes feasible even for cases when there are hundreds of thousands of features.

### 6.6 Model fitting

All models were fitted on the log2 values of the MIC, which is the natural scale for the MIC measurement. Later the predicted values were converted back to the MIC units.

### 6.7 Study pipeline

The pipeline of this study is shown in Figure 2. First, the raw reads were converted to a numerical table indicating mutations and acquired resistance encoding genes. In the second step, a random forest model is fitted on the train data via leave-one-country cross-validation. Features importances were averaged over each fold. Then the highest-ranking features were kept which significantly reduced the dimensionality of the data. Using this low dimensional training data a random forest model and a linear regression were fitted. Regression models were preferred because they can have a precise susceptibility level estimation instead of classification, which has only a binary susceptibility outcome. For fitting the models always the log2 MIC values were used as a natural scale for the MIC measurements. At the last step, the performance of the models was evaluated on the unseen test data using the same restricted feature set.

**FIG 1.**
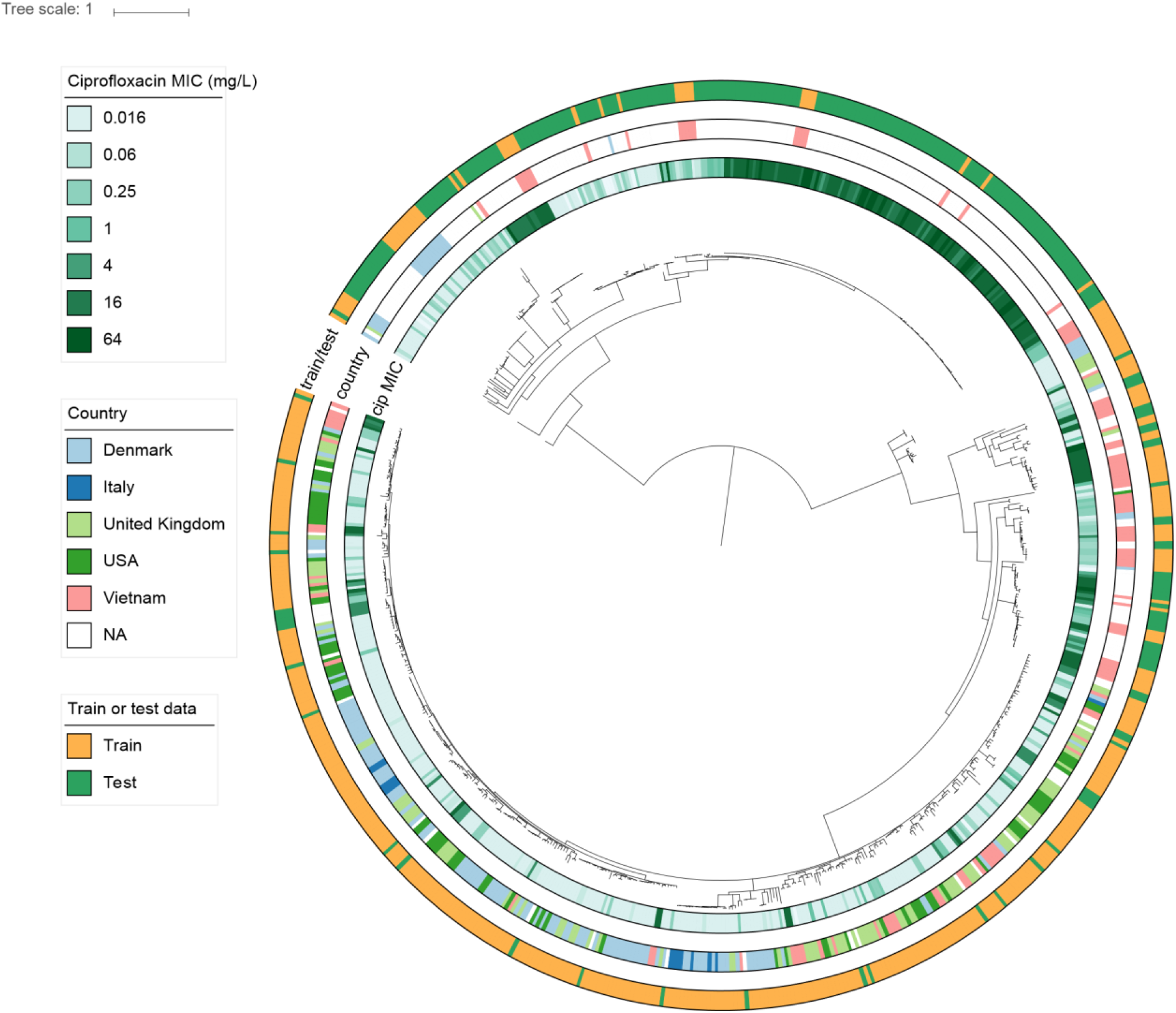
Midpoint-rooted phylogenetic tree of the 704 *E. coli* samples that had ciprofloxacin MIC measurement. It is clearly visible that the test data is clustered separately from the training data suggesting the generalization power of our model. Nodes with lower than 80% bootstrap support are collapsed.

**FIG 2.**
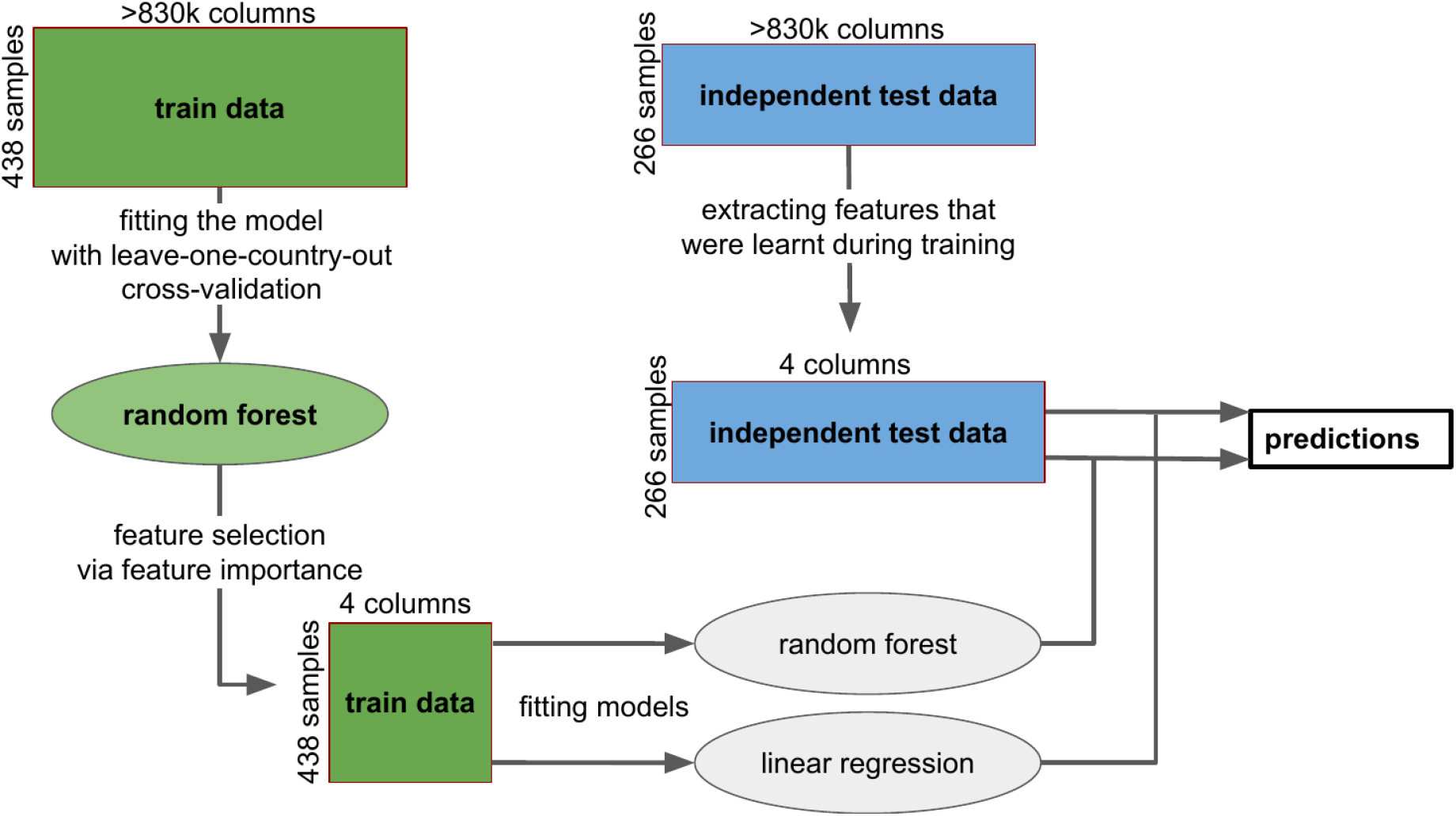
Workflow of the study. First, a random forest model was fitted to the training data with leave-one-country-out validation. Feature importances of the fitted models are averaged over all the folds and the four best features are kept. Then the random forest model and a linear regression model were fitted on all the training samples using only the four best features. And model performances are tested using the independent test dataset.

## 7. Results

Our dataset consists of a phylogenetically diverse collection of *E. coli* strains (Fig. 1). Strains in the test and train group are present throughout the whole phylogeny, although the groups are present predominantly in different parts of the phylogeny.

We trained a machine learning model using genome-wide mutation profiles alongside the ResFinder-based profiles of acquired resistance genes. We ranked the predictors proposed by the model itself, see Table S1. The model performed with high accuracy on the training set leave-one-country-out cross-validation using four predictors, see Fig S1. The addition of more features did not improve the cross-validation results, and therefore we kept only the first four, allowing for a simple and understandable model.

Using these four predictors, 265 out of the 266 test data samples were correctly classified by our models at susceptible/non-susceptible level, and more than 92% of the corresponding MIC values were correctly predicted within a four-fold dilution, see Table 2.

**TABLE 2.** Number of features, R^2^ score, Pearson correlation (R), Major Error (ME), Very Major Error (VME), area under the receiver operating curve (AUC), Accuracy within a two/four-fold dilution (ACC-2, ACC-4) and Mean Absolute Fold Error (MAFE) on the unseen test data. For the AUC, ME, VME the data was binarized using 1 mg/L threshold. The number of features were selected according to the performance using leave-one-country-out validation on the training data, see Fig.S1

| model | N_feat | <sup>+</sup>R<sup>2</sup> | <sup>+</sup>R | <sup>#</sup>ME | <sup>#</sup>VME | AUC | ACC-2 | ACC-4 | <sup>#</sup>MAFE |
| --- | --- | --- | --- | --- | --- | --- | --- | --- | --- |
| random forest | 4 | 0.932 | 0.966 | 1 | 0 | 1.000 | 0.654 | 0.944 | 0.891 |
| random forest | 15 | 0.890 | 0.944 | 4 | 0 | 0.998 | 0.684 | 0.891 | 1.007 |
| linear regression | 4 | 0.914 | 0.957 | 1 | 0 | 1.000 | 0.654 | 0.921 | 0.998 |

\* number of samples
<sup>+</sup>calculated on the log2 values
<sup>#</sup>the lower the better

These 4 predictors are the following:

1. *gyrA* mutation at amino acid #87
2. *gyrA* mutation at amino acid #83
3. *parC* mutation at amino acid #80
4. presence of any *qnrS* gene

All of the predictors above are binary (presence/absence) therefore there are 2^4^ = 16 different possible prediction for any sample based on these features, see Table S2. A linear regression model fitted on the log2 values of the MIC measurements could achieve similar performance as a more complex random forest model, see Figure 3. Linear regression is preferred due to its simplistic nature. Having a random forest regressor with hundreds of decision trees and thousands of genomic features as predictors it is difficult to understand why the model made that particular prediction, leaving doubts of its clinical usefulness.

**FIG 3.**
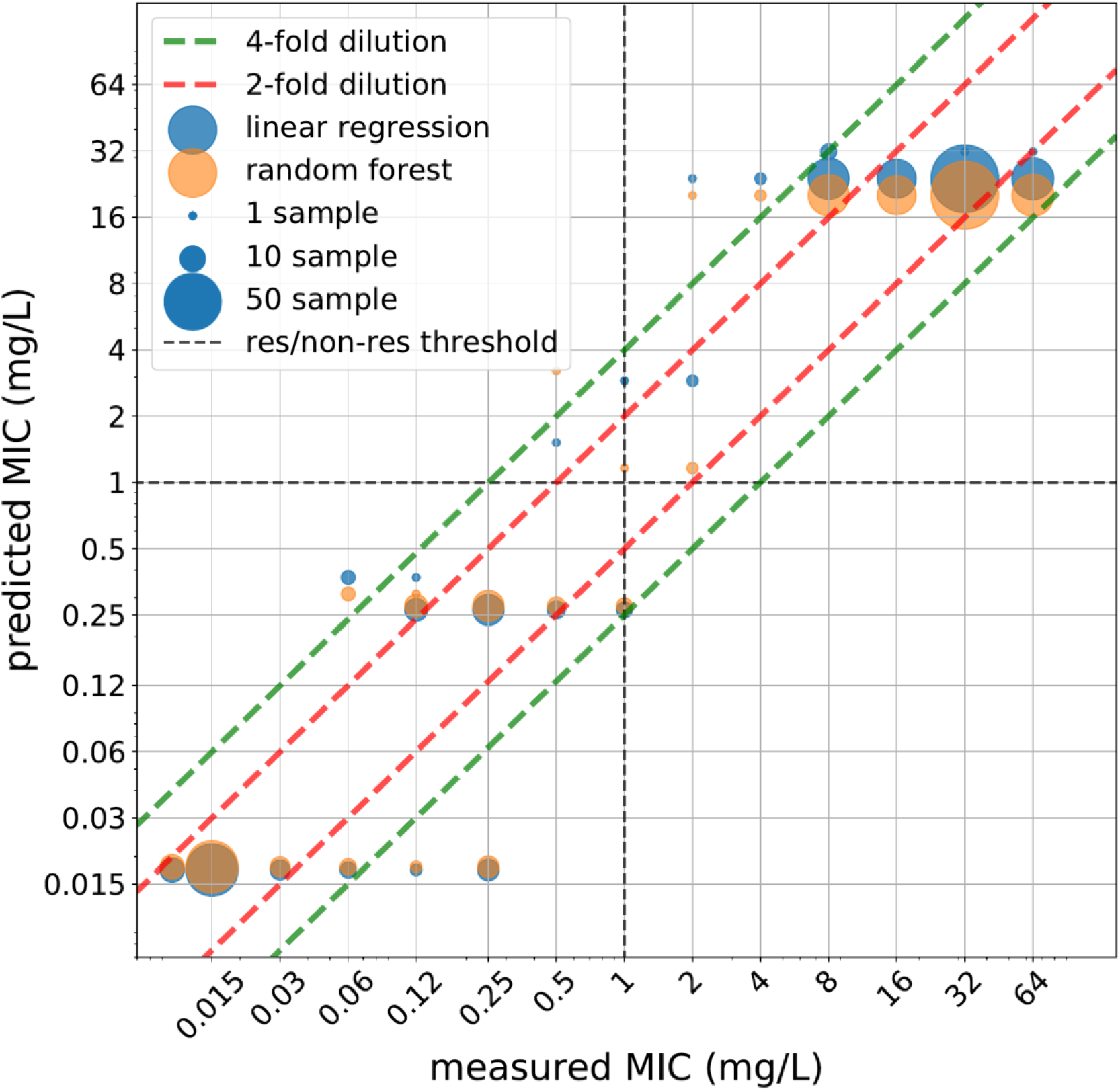
Prediction on the unseen test set was generated via random forest and linear regression model using the best four predictors. It can be clearly seen that the two models do not differ much in terms of predicted values.

## 8. Discussion

Here, we present a novel method for predicting ciprofloxacin resistance for *E. coli*. With minimal prior knowledge of relevant resistance determinants (that is mainly the use of ResFinder which includes genes known to encode low-level resistance to ciprofloxacin in *E. coli*) and a data-driven approach, we managed to create a machine learning model that was not only accurately predicting the susceptible/non-susceptible labels but also accurately predicting at MIC level. Additionally, the highlighted features of our approach could be narrowed down to four biologically understandable features, making the method simpler and therefore applicable to clinical microbiology practice.

It was previously shown that accurate ciprofloxacin resistant/susceptible binary prediction is possible for *E. coli* (17) (12) (3). For some other bacteria-antibiotic combinations even MIC level predictions were performed (19), (20), (18), (21). This study goes beyond by not only predicting MIC level ciprofloxacin resistance for *E. coli,* but also highlighting the underlying reasoning behind the predictions. Furthermore, this study is one of very few that includes the presence or absence of genes located on mobile genetic elements (MGEs), in combination with chromosomal point mutations, in the machine learning algorithm. This is a crucial step since particularly in Gram-negative microorganisms such as *E. coli*, AMR is often encoded by genomic determinants located on MGE, or a combination of chromosomal and MGE encoded determinants, as is demonstrated in our study for ciprofloxacin. In addition, this study used data from different countries and regions thus ensuring potential variation in determinants that may contribute to ciprofloxacin resistance are represented in the data set.

Notably, a linear regression model based on only the four most important features of the random forest model performed nearly as well as the full model. These features comprise two *gyrA* mutations, one *parC* mutation and the presence of any *qnrS* gene. All features have been associated with ciprofloxacin resistance before (23). In addition, the presence of a single determinant versus combinations of multiple of the four determinants predicted MIC ranges that were comparable to those observed experimentally and in clinical isolates (23). For example, the single presence of a *qnr* gene predicted a relatively low MIC but the combination of a *qnr* gene with a single mutation in *gyrA* increased the predicted MIC substantially (Table S2). Our results indicate that for prediction of ciprofloxacin susceptibility on the basis of whole-genome sequencing in *E. coli,* the analysis could be limited to only these four determinants.

It is worthy to note that the model was trained on all possible mutations present in the data set, not only known mutations or acquired resistance genes from a curated database. Therefore the model could potentially discover novel currently unknown mutation-based resistance encoding mechanisms which may be located in genes that are or are not yet known to contribute to resistance. For *E. coli* the ciprofloxacin resistance determinants that were predicted in our machine learning approach have been experimentally verified, but for other antibiotics, our approach could detect novel genomic variants associated with resistance.

Our study also has some limitations, which mostly pertain to the dataset. For strains with measured MICs in the range of 8-64 mg/L, our model performs worse than for strains with lower MICs. This is most likely due to the fact that the majority of resistant strains in our training data have an MIC of 32 mg/L, with only very few other resistant MICs. This hampers accurate prediction of MIC for more resistant *E. coli*. Additionally, our dataset is not yet diverse and complete enough to be applied on a wide scale. This is a common problem for many studies aiming to predict AMR from WGS data. Solving this would require continuous updating of databases and an adequate database structure, the latter we have addressed previously (24).

In conclusion, we report a novel machine learning approach for a quantitative prediction of antibiotic resistance, which we applied for prediction of ciprofloxacin resistance in *E. coli*. In combination with continuous data base improvements, our approach could allow machine learning methods to enter routine clinical diagnostic and epidemiological practices to continuously improve predictions.

## 9. Author statements

### 9.1 Authors and contributors

All authors contributed to the conceptualization of the study. B.A.P., B.C.L.P. and C.S. wrote the manuscript, S.M., B.C.L.P., R.S.H., O.L., C.S. collected the data from various public data sources. B.A.P., E.G., D.A.A. performed machine learning modeling. The COMPARE ML-AMR group contributed data and technical help. All authors contributed to the study with insightful discussion. All authors reviewed the manuscript.

## 9.4 Acknowledgements

The COMPARE ML-AMR group: S. Matamoros^1^; V. Janes^1^; R. S. Hendriksen^2^; O. Lund^2^; P. Clausen^2^; Frank M. Aarestrup^2^; Marion Koopmans^6^; B. Pataki^3,4^; D. Visontai^3,4^; J. Stéger^3,4^; JM. Szalai-Gindl^3,4^; I. Csabai^3,4^; N. Pakseresht^5^; M. Rossello^5^; N. Silvester^5^; C. Amid^5^; G. Cochrane^5^; C. Schultsz^1,7^, F. Pradel^8^; E. Westeel^8^; S. Fuchs^9^; S. Malhotra Kumar^10^; B. Britto Xavier^10^; M. Nguyen Ngoc^10^; D. Remondini^11^; E. Giampieri^11^; F. Pasquali^12^; L. Petrovska^13^; D. Ajayi^13^; E. M. Nielsen^14^; N. V. Trung^15^; N. T. Hoa^15^; Yoshikazu Ishii^16^; Kotaro Aoki^16^; P. McDermott^17^.

^1^: Amsterdam UMC, University of Amsterdam, Department of Medical Microbiology, Amsterdam, The Netherlands.

^2^: National Food Institute, Technical University of Denmark, Lyngby, Denmark.

^3^: Department of Physics of Complex Systems, ELTE Eötvös Loránd University, Budapest, Hungary.

^4^: Department of Computational Sciences, Wigner Research Centre for Physics of the HAS, Budapest, Hungary.

^5^: European Molecular Biology Laboratory, European Bioinformatics Institute, Wellcome Genome Campus, Hinxton, Cambridge CB10 1SD, UK.

^6^: Department of Viroscience, Erasmus University Medical Center, Rotterdam, The Netherlands.

^7^: Amsterdam UMC, University of Amsterdam, Department of Global Health, Amsterdam Institute for Global Health and Development, Amsterdam, The Netherlands.

^8^: Fondation Mérieux, Lyon, France.

^9^: Department of Infectious Diseases, Robert Koch Institut, Berlin, Germany.

^10^: Department of Medical Microbiology, Vaccine & Infectious Disease Institute, Antwerp University, University Hospital Antwerp, Antwerp, Belgium.

^11^: Department of Physics and Astronomy (DIFA), University of Bologna, Bologna, Italy.

^12^: Department of Agricultural and Food Sciences (DISTAL), University of Bologna, Bologna, Italy.

^13^: Animal and Plant Health Agency, Addlestone, Surrey, United Kingdom.

^14^: Statens Serum Institut, Denmark.

^15^: Oxford University Clinical Research Unit, Centre for Tropical Medicine, Ho Chi Minh City, Vietnam.

^16^: Department of Microbiology and Infectious Diseases, Faculty of Medicine, Toho University School of Medicine, 5-21-16 Omorinishi, Ota-ku, Tokyo 143-8540, Japan.

^17^: Food and Drug Administration, Center for Veterinary Medicine, Office of Research, Laurel, MD, USA.

## 11. Supplemental material file list

- **TABLE S1** shows the feature importances over the different leave-one-country-out folds
- **TABLE S2** shows the parameters of the fitted linear regression and the 2^4^ = 16 possible predicted values based on the 4 features.
- **FIGURE S1** shows the leave-one-country-out cross-validation R^2^ results based on the number of features.
- **FIGURE S2** shows the results of the models for an additional 100 *E. coli* genomes from Bangladesh. These genomes had only disk diffusion test measurements, which is why they were not discussed in the paper.
- **FIGURE S3** shows data quality control checks for the dataset.

## 12. Data bibliography

All the used data is publicly available from the SRA EBI database. All used files are listed at https://github.com/patbaa/AMR_ciprofloxacin/blob/master/meta.tsv with URLs provided. Isolates with accession numbers and MIC measurements are also available at https://github.com/patbaa/AMR_ciprofloxacin/blob/master/supplementary_meta_table.csv.

---

**FIG S1.**
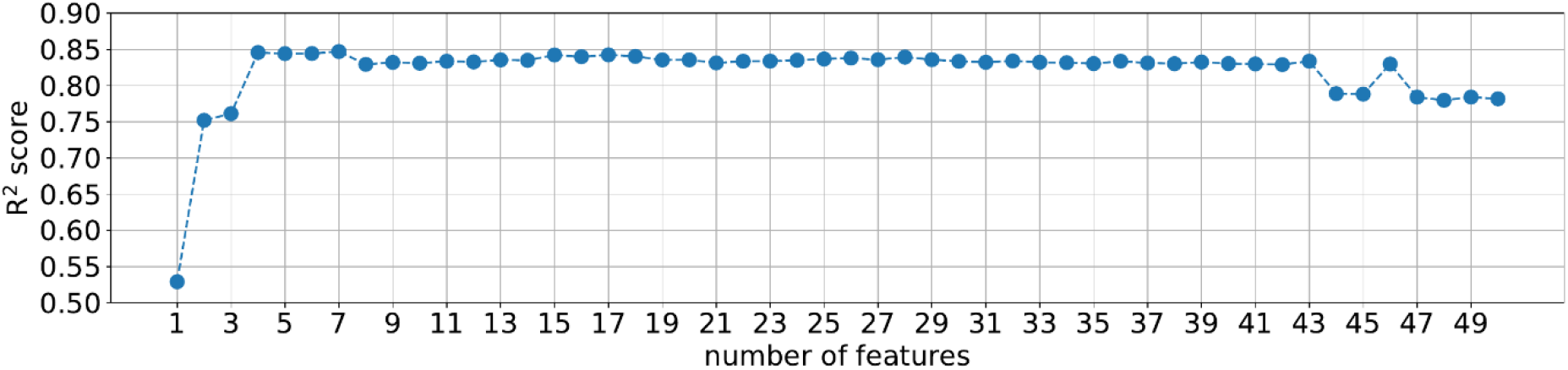
R squared score calculated on the training set using random forest model. The features were ranked based on Table S1 and iteratively a random forest model was fitted on the training set with leave-one-country-out validation.

**FIG S2.**
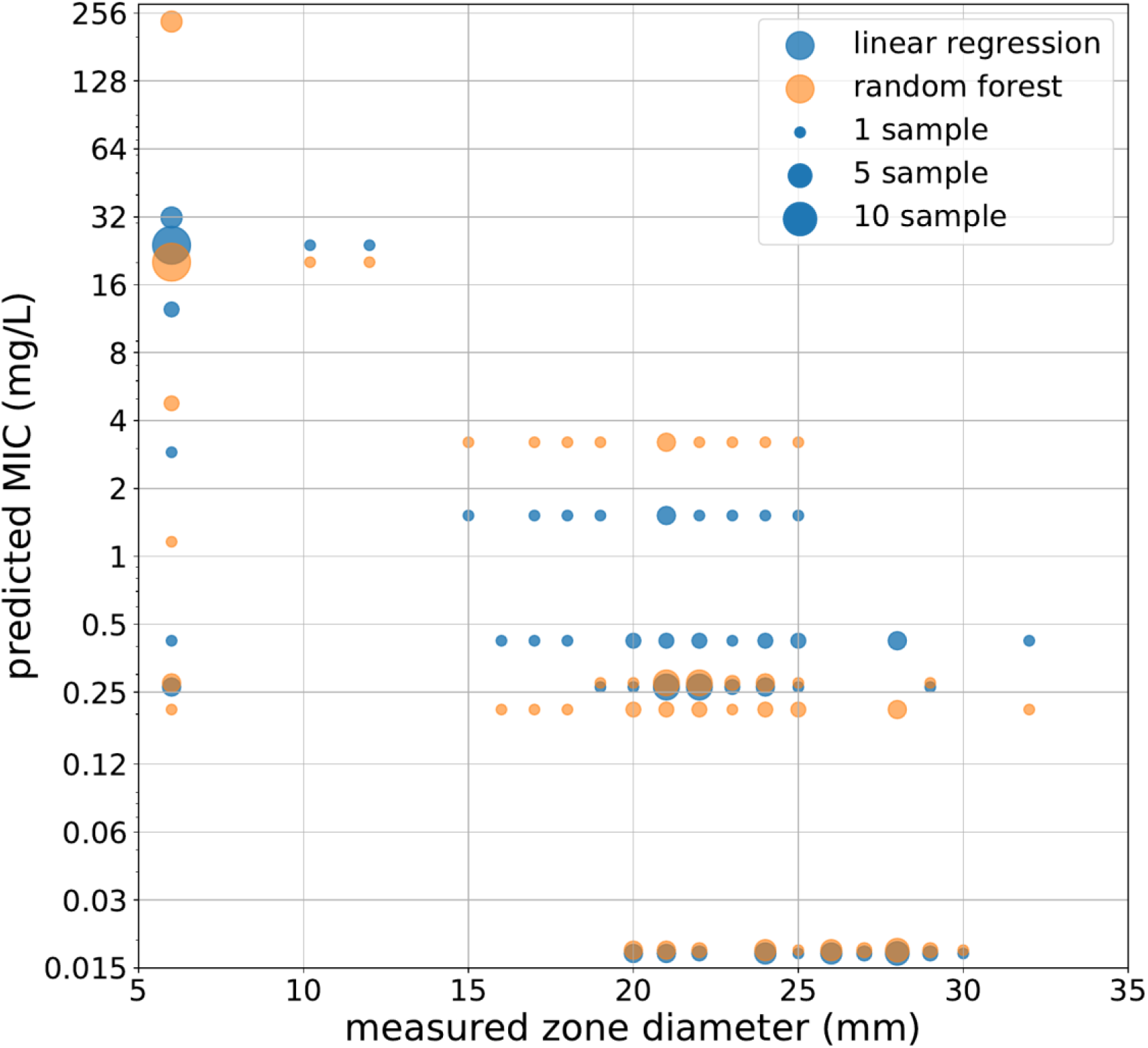
Prediction for samples that had only disk diffusion test measurement. As the larger zone diameter corresponds to smaller MIC values, a negative correlation is desirable on this plot. The same models were used with 4 predictors as it was used for the test set.

**FIG S3.**
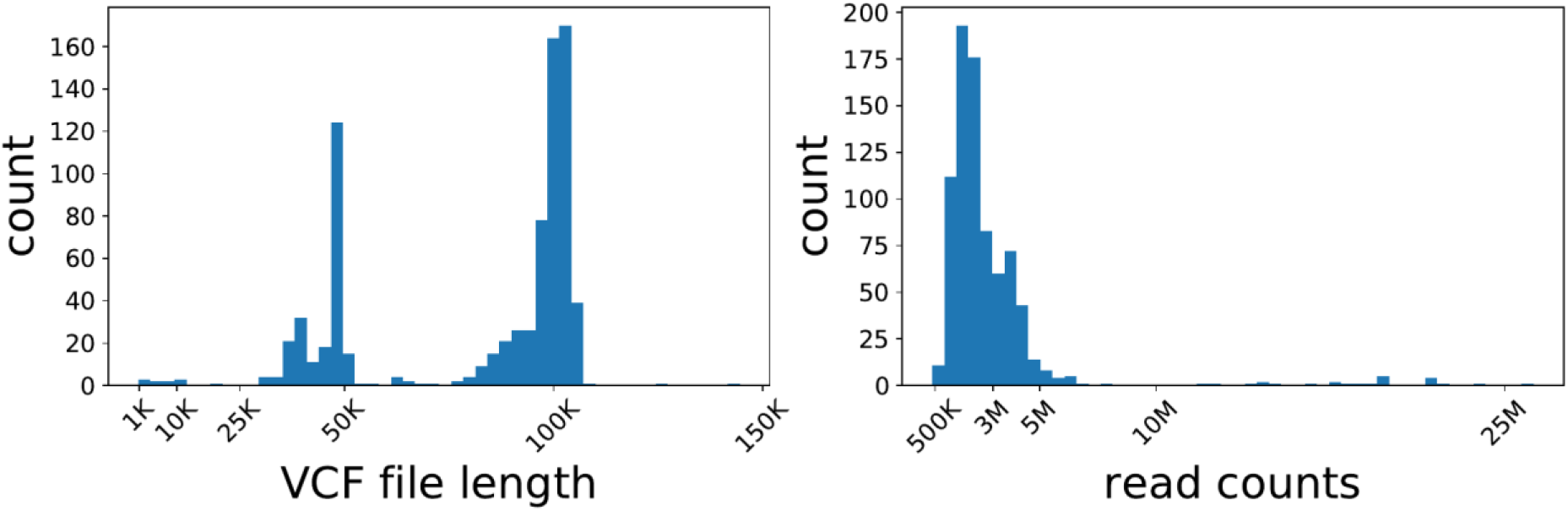
VCF file length distribution and the number of raw reads in the collected dataset.

**TABLE S1.** Feature importances of the fitted random forest models. Random forest model was fitted on the training data using leave-one-country-out validation. Each entry shows the feature importance for the given feature for the validation step when the isolates from the given country were not used to train the model. Sorted by the sum of the feature importances. The features are following the gene # amino acid position naming where possible. For the mutations where there were no genes associated, the naming is chromosome name _ position. For the features coming from ResFinder, the ResFinder naming was kept. has_qnr and has_qnrS are binary features describing if the isolate had any qnr/qnrS entry in the ResFinder results.

| feature | Denmark | Italy | USA | UK | Vietnam | sum |
| --- | --- | --- | --- | --- | --- | --- |
| gyrA#87 | 0.555 | 0.567 | 0.570 | 0.558 | 0.010 | 2.260 |
| gyrA#83 | 0.114 | 0.148 | 0.107 | 0.141 | 0.690 | 1.200 |
| parC#80 | 0.181 | 0.151 | 0.199 | 0.172 | 0.001 | 0.705 |
| has_qnrS | 0.042 | 0.039 | 0.019 | 0.037 | 0.004 | 0.140 |
| qnrS1_1_AB187515 | 0.012 | 0.007 | 0.017 | 0.004 | 0.000 | 0.041 |
| blaCTX-M-55_1_DQ810789 | 0.002 | 0.009 | 0.009 | 0.013 | 0.000 | 0.033 |
| blaVIM-48_1_KY362199 | 0.004 | 0.001 | 0.016 | 0.002 | 0.000 | 0.022 |
| CP009072.1_3517597 | 0.000 | 0.000 | 0.000 | 0.000 | 0.013 | 0.014 |
| CP009072.1_1734215 | 0.000 | 0.000 | 0.000 | 0.000 | 0.008 | 0.009 |
| blaCTX-M-14_1_AF252622 | 0.000 | 0.002 | 0.002 | 0.003 | 0.000 | 0.008 |
| CP009072.1_3517591 | 0.000 | 0.000 | 0.000 | 0.000 | 0.007 | 0.007 |
| CP009072.1_113480 | 0.000 | 0.000 | 0.000 | 0.000 | 0.005 | 0.006 |
| CP009072.1_1205372 | 0.003 | 0.002 | 0.001 | 0.000 | 0.000 | 0.006 |
| has_qnr | 0.005 | 0.000 | 0.000 | 0.000 | 0.000 | 0.005 |
| CP009072.1_459777 | 0.001 | 0.002 | 0.000 | 0.002 | 0.000 | 0.005 |

**TABLE S2.**
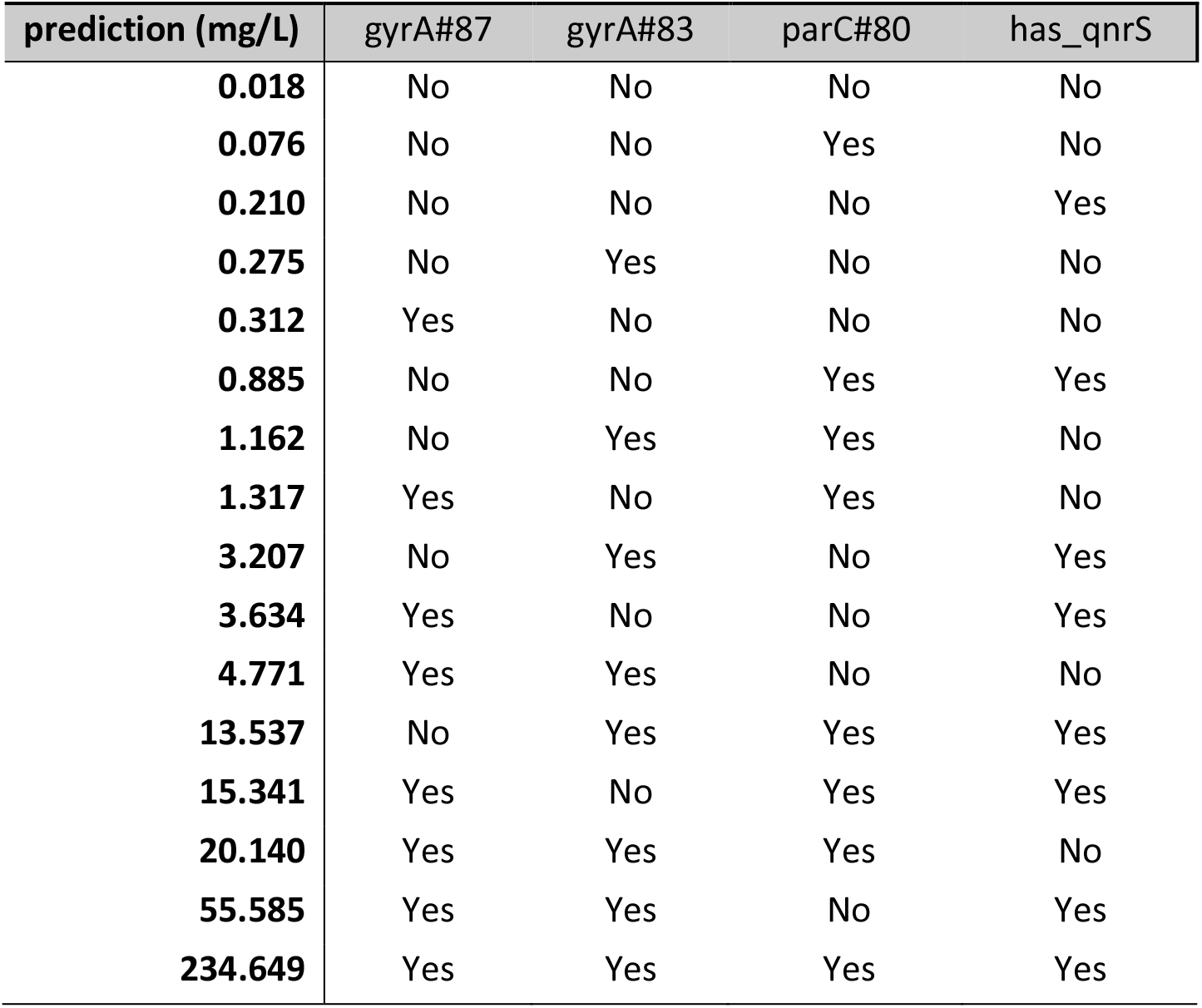
Parameters of the fitted linear regression model. The interception is −5.796, and the parameters associated with gyrA#87, gyrA#83, parC#80, has_qnrS are 4.116, 3.935, 2.078 and 3.542. Prediction is calculated as 2 to the power of the sum of interception and the present mutation/genes.

| prediction (mg/L) | gyrA#87 | gyrA#83 | parC#80 | has_qnrS |
| --- | --- | --- | --- | --- |
| 0.018 | No | No | No | No |
| 0.076 | No | No | Yes | No |
| 0.210 | No | No | No | Yes |
| 0.275 | No | Yes | No | No |
| 0.312 | Yes | No | No | No |
| 0.885 | No | No | Yes | Yes |
| 1.162 | No | Yes | Yes | No |
| 1.317 | Yes | No | Yes | No |
| 3.207 | No | Yes | No | Yes |
| 3.634 | Yes | No | No | Yes |
| 4.771 | Yes | Yes | No | No |
| 13.537 | No | Yes | Yes | Yes |
| 15.341 | Yes | No | Yes | Yes |
| 20.140 | Yes | Yes | Yes | No |
| 55.585 | Yes | Yes | No | Yes |
| 234.649 | Yes | Yes | Yes | Yes |

